# Nucleus Basalis links sensory stimuli with delayed reinforcement to support learning

**DOI:** 10.1101/663211

**Authors:** W. Guo, D.B. Polley

## Abstract

Linking stimuli with delayed reinforcement requires neural circuits that can bridge extended temporal gaps. Auditory cortex (ACx) circuits reorganize to support auditory fear learning, but only when afferent sensory inputs temporally overlap with cholinergic reinforcement signals. Here we show that mouse ACx neurons rapidly reorganize to support learning, even when sensory and reinforcement cues are separated by a long gap. We found that cholinergic basal forebrain neurons bypass the temporal delay through multiplexed, short-latency encoding of sensory and reinforcement cues. At the initiation of learning, cholinergic neurons in Nucleus Basalis increase responses to conditioned sound frequencies and increase functional connectivity with ACx. By rapidly scaling up responses to sounds that predict reinforcement, cholinergic inputs jump the gap to align with bottom-up sensory traces and support associative cortical plasticity.

## Introduction

Animals rapidly learn to fear a neutral sound stimulus (conditioned stimulus, CS) that overlaps in time with an aversive stimulus (unconditioned stimulus, US). Behavioral evidence of learning can be indexed by conditioned behavioral changes such as freezing or escape elicited by a CS that has been paired with the US (CS+), but not by a CS that fails to predict US onset (CS-). Auditory fear learning reflects a plasticity at sites of CS and US convergence, including the amygdala and auditory cortex (ACx) (Letzkus et al., 2015; McGann, 2015; Weinberger, 2004). Aversive stimuli trigger the phasic release of acetylcholine (ACh) in ACx via projections from the basal forebrain, which acts through local inhibitory microcircuits to modulate the excitability of cortical pyramidal neurons (Froemke et al., 2007; Kuchibhotla et al., 2017; Letzkus et al., 2011; Nelson and Mooney, 2016; Takesian et al., 2018; Urban-Ciecko et al., 2018). During brief windows of disinhibition, the afferent CS+ trace elicits enhanced spiking of cortical pyramidal neurons (Froemke et al., 2007; Letzkus et al., 2011, 2015; Pi et al., 2013) and induces a persistent and selective potentiation of the CS+ frequency that can be indexed through increased excitatory synaptic strength (Froemke et al., 2007, 2013), increased spiking (Bakin and Weinberger, 1990, 1996; Froemke et al., 2013; Quirk et al., 1997), over-representation of the CS+ frequency in ACx tonotopic maps of sound frequency (Kilgard and Merzenich, 1998) and enhanced perceptual awareness of the CS+ sound (Aizenberg et al., 2015; Froemke et al., 2013; Reed et al., 2011).

## Results

Explanatory models for local circuit changes that enable associative learning emphasize the importance of the temporal “handshake” arising from the convergence of the CS-evoked sensory trace and the US-evoked cholinergic trace (**Fig. 1A**) (Chubykin et al., 2013; Letzkus et al., 2011, 2015; Metherate and Ashe, 1991). To test the prediction that phasic ACh release from basal forebrain neurons is sufficient to enhance ACx spiking responses to co-occurring sound frequencies, we optogenetically activated cholinergic axon terminals in the ACx by pulsing blue light on the cortical surface of awake, head-fixed ChAT-Cre mice crossed to a Cre-dependent channelrhodopsin2 reporter line (**Fig. 1B**). Extracellular recordings were made from the ACx of awake, head-fixed mice with multi-channel silicon probes (**Fig. 1C**). Putative pyramidal neurons with regular spiking waveforms (**Fig. S1A**) were held for approximately 80 minutes, during which time auditory receptive fields were characterized with tone pips of varying frequency (Test blocks, Fig. 1C) interspersed with six conditioning blocks of 10 trials each. In support of the phasic ACh release model, pairing narrowband noise bursts (NBN) with simultaneous optogenetic activation of ACx cholinergic axon terminals resulted in selectively enhanced spiking at the paired frequency (Bakin and Weinberger, 1996; Froemke et al., 2013) (n = 58 units, ANOVA main effect for conditioning stage F = 6.64, p < 0.01, one-sample t-tests against a population mean of zero, p < 0.025 for all conditioning stages, **Fig. 1D-G**).

**Fig. 1.**
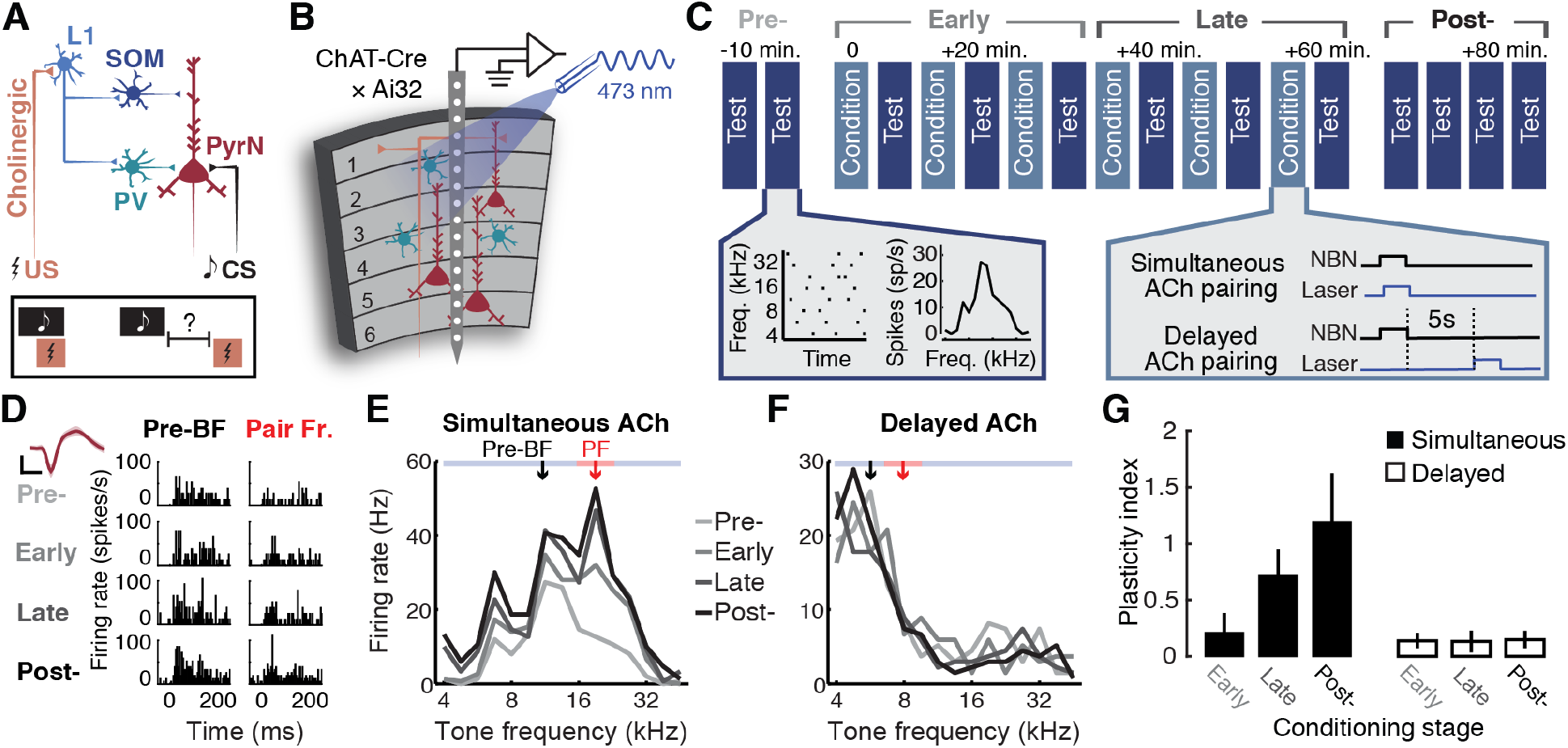
Cholinergic terminal activation in auditory cortex induces associative receptive field plasticity, but only when temporally overlapping with the CS. (***A***) Cartoon of ACx circuitry for plasticity and learning when auditory conditioned stimuli (CS) are paired with aversive unconditioned stimuli (US). L1 = layer 1; SOM = somatostatin-expressing; PV = parvalbumin. (***B***) Schematic of strategy to induce ACx receptive field plasticity with local optogenetic activation of cholinergic terminals in ChAT-Cre × Ai32 mice. (***C***) Optogenetic pairing and testing protocol for awake head-fixed ACx recording experiments. NBN = narrow band noise. (***D***) Spike PSTHs from a regular spiking unit at the pre-pairing best frequency (BF) and the frequency paired with cholinergic terminal activation (Pair Fr. or PF). Inset: single unit spike waveform, where shading = SEM. Scale bars = 50 μV and 0.4ms. (***E***) Frequency response functions for the same RS unit across conditioning stages. The fold change in normalized spike rate at frequencies far away from the PF (> ± 0.25 octaves, blue) is subtracted from frequencies near the PF (≤ 0.25 octaves, red) to compute the Plasticity Index, where values greater than 0 indicate enhanced firing rates near the paired frequency. (***F***) Frequency response functions from an example unit where a 5s delay separated the offset of sound stimuli and cholinergic axon stimulation. (***G***) mean ± SEM plasticity index across conditioning stages (n =58/20 units for simultaneous/delayed ACh pairing). All statistical tests are adjusted for inflated false discovery using the Holm–Bonferroni correction for multiple comparisons.

By contrast to the effect of simultaneous pairing of NBN and cholinergic axon activation, Receptive field remodeling was not observed when laser onset was delayed by 5s from the cessation of the CS+ stimulus (n = 20, ANOVA main effect for conditioning stage F = 0.58, p = 0.44, one-sample t-tests against a population mean of zero, p > 0.07 for all conditioning stages, **Fig. 1E-G**). In the ACx, electrophysiological signatures of brief sounds fade away within a few hundred milliseconds following sound offset (**Fig. S1B-C**). Therefore, CS-specific plasticity would not be expected to occur when the cholinergic surge arrived in ACx long after the sensory trace had expired (Metherate and Ashe, 1991). Yet behaviorally, associative learning can occur even when the US begins several seconds or more after the CS ends (Garcia et al., 1966; Pavlov, 1932). To determine whether auditory fear learning could also bridge an extended temporal gap, we conditioned mice over five consecutive days either with a 16 kHz NBN CS+ followed 5s later by a foot shock (N=4) or a no-shock control condition (N=4) (**Fig. 2A**). On day 6, both groups of mice were introduced to a novel sensory context and presented with CS+ and CS- sound frequencies. We found that mice conditioned with a 5s gap between the CS and US exhibited associative learning that appeared virtually identical to previous descriptions with temporally overlapping sounds and foot shocks (Letzkus et al., 2011) (ANOVA Group × Frequency interaction, F = 32.3, p < 0.0001; **Fig. 2B**).

**Fig. 2.**
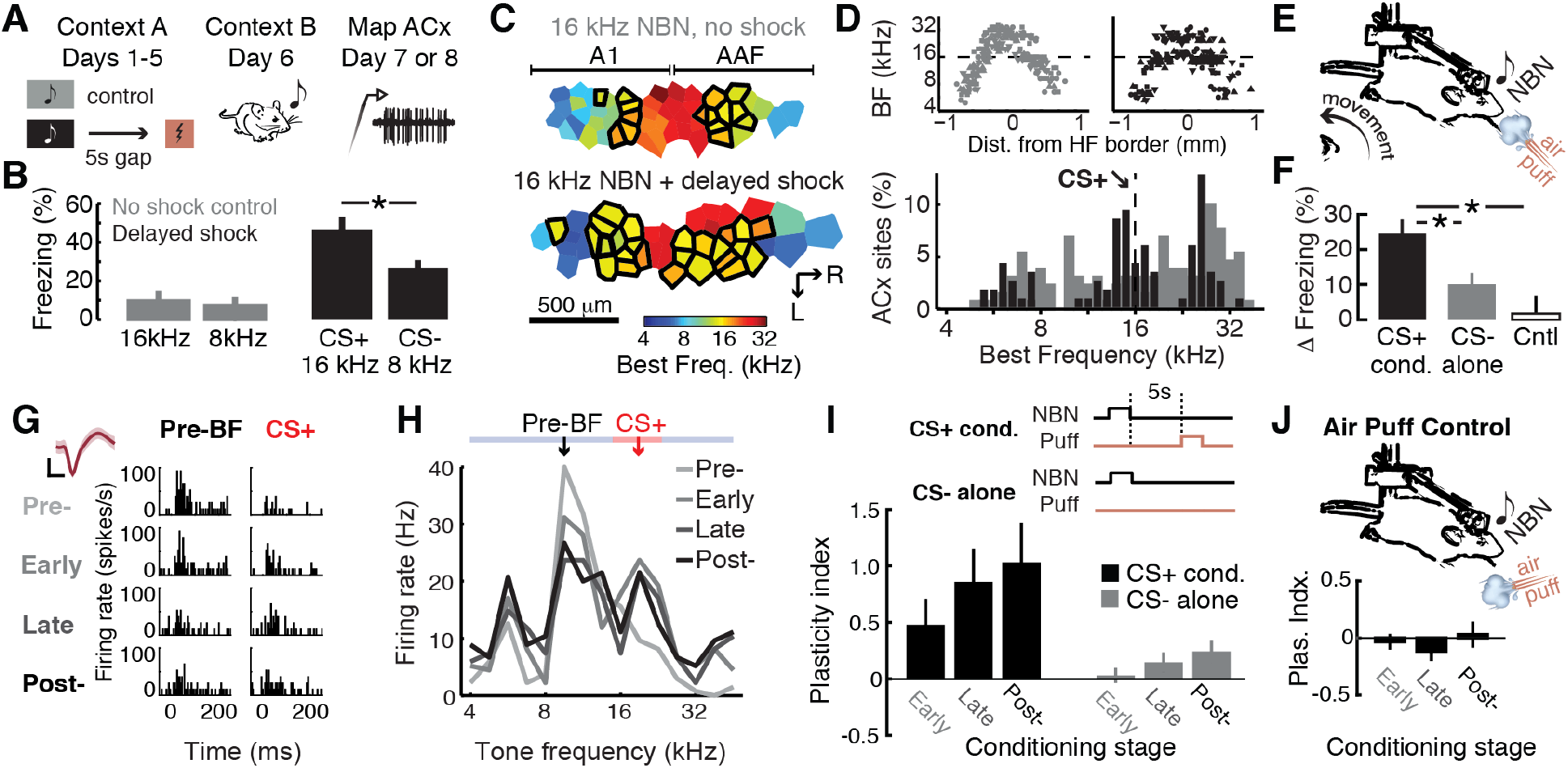
Rapid emergence of receptive field plasticity and associative learning despite a long delay separating the CS and aversive reinforcement. (***A-B***) Conditioned freezing behavior to CS+ tone is observed when a 5s delay separates the CS and US. Asterisk indicates a significant Group × Frequency interaction term. Bar plots show mean ± SEM. (***C***) Tonotopic maps of best frequency (BF) are reconstructed from microelectrode recordings of multiunit tonal receptive fields from the primary auditory cortex (A1) and anterior auditory field (AAF) (Control, n = 118/4; Delayed shock conditioning, n = 130/4 [recordings sites/mice]) of the mice behaviorally characterized in (*B*). (***D***) Distortions in the caudal-to-rostral BF mirror reversal gradient (*top*) over-represent the 16 kHz CS+ frequency, as shown in histograms of BF distributions in tone-alone (gray) and tone + delayed shock (black) groups. (***E***) Preparation to study the emergence of plasticity and conditioned freezing in head-fixed mice. (***F*** Freezing behavior on the rotary treadmill. Asterisks indicate significant differences between CS+ conditioning, in which narrow band noise (NBN) precedes air puffs to the face by 5s (N=16), CS- alone conditioning, where NBN is presented without air puffs and a control where NBN precedes air puffs that are aimed away from the face (N=5). Bar plots show mean ± SEM. (***G***) Tone-evoked PSTH from a representative regular spiking unit across conditioning stages. Sale bars are 20 μV and 0.4 ms. (***G***) Frequency receptive field from the same unit across conditioning stages. The fold change in normalized spike rate for frequencies far from the CS (blue) is subtracted from that at or near the conditioned stimulus (red) to compute the Plasticity Index, where values greater than 0 denote CS-specific frequency enhancement. (***I***) Mean ± SEM Plasticity Index across conditioning stages (n =24/58 units for CS+ conditioning/CS- alone). (***J***) Experiments where CS+ conditioning is performed with the air puff nozzle pointed away from the face (n =20). All statistical tests are adjusted for inflated false discovery using the Holm–Bonferroni correction for multiple comparisons.

Because the transient ACh “handshake” model for ACtx plasticity and auditory fear learning has only been studied with protocols where CS and US traces overlap in time, one possibility is that ACx plasticity may not contribute to auditory fear learning when the CS and US are separated by longer temporal gaps. Instead, the critical locus of plasticity could shift to other brain areas such as the hippocampus, entorhinal cortex or medial prefrontal cortex (Raybuck and Lattal, 2014). We addressed this possibility by quantifying tonotopic maps of best frequency (BF) from multiunit recordings in the primary auditory cortex (A1) and the anterior auditory field (AAF) from the same eight mice that underwent behavioral testing (**Fig. 2C**). We found that tonotopic receptive field maps were distorted in mice conditioned with a 5s CS-US delay (n = 130 recording sites), such the 16 kHz CS+ frequency was over-represented at the expense of neighboring frequencies, as compared to the smooth BF gradients measured in mice passively exposed to the CS+ and CS- sounds (n = 118 recording sites; 2-sample K-S test, p < 0.05; **Fig. 2D**).

These findings demonstrate that associative auditory fear learning can occur with an extended US delay, but leaves open the questions of where, when and how associative plasticity is initially formed during the early stages of learning. To track the parallel emergence of associative learning and cortical plasticity, we returned to the awake head-fixed single unit recording preparation. Mice either underwent CS+ conditioning (NBN followed 5s later by an air puff to the face) or CS- alone (NBN without air puff). Mice could walk freely on a rotating disk, providing us with the means to measure NBN-elicited freezing during conditioning blocks while head-fixed (**Fig. 2E**). We found that cortical plasticity and auditory learning emerged shortly after the start of conditioning, despite the 5s gap between the CS and US. NBN-triggered freezing behavior was significantly increased in CS+ Conditioning blocks compared to either CS- alone or a control in which air puffs were presented at the 5s delay but were directed away from the face (ANOVA, main effect for Group, F = 6.51, p < 0.005; pairwise comparisons, p < 0.05 and 0.01, respectively, after Holm–Bonferroni correction for multiple comparisons; **Fig. 2F**). Importantly, receptive field plasticity induced by paring sound and punishment with a 5s delay was comparable to changes induced by simultaneous pairing of sound and local cholinergic axon activation. CS+-specific increases in spiking grew steadily during the early and late stages of conditioning and remained elevated after conditioning ended (CS+ conditioning, ANOVA, F = 8.39, p < 0.005; **Fig. 2G-H**). CS-specific changes in frequency response functions were significantly greater in CS+ conditioning than CS- conditioning, where NBN bursts were not paired with air puffs (n = 24/58 for CS+/CS-, Wilcoxon Rank Sum, p < 0.01; **Fig. 2I**). Likewise, CS-specific changes were not observed in a control conditioning group where air puffs were directed away from the face (One-way ANOVA, F = 0.84, p = 0.36; n = 20; **Fig. 2J**).

These findings were puzzling. On the one hand, we observed selective and persistent enhancement of sound frequencies paired with simultaneous cholinergic terminal activation, but not when ACh terminal activation lagged sound offset by 5s (Fig. 1g). On the other hand, selective and persistent frequency enhancement was observed despite the 5s gap when an aversive US was used rather than local ACx cholinergic activation. This lead us to question the assumption that cholinergic inputs from the basal forebrain were only elicited by the US. If instead, cholinergic basal forebrain inputs were activated by the CS as well as the US during learning, the cholinergic input would functionally bypass the 5s gap and temporally coincide with the afferent sensory trace to establish the CS-US temporal handshake.

To test this hypothesis, we recorded from optogenetically identified cholinergic basal forebrain neurons that project to the ACx (ChACx). To identify the highest density of ChACx neurons, we first performed an anatomy experiment in which fluorescent microbeads were injected into A1 of ChAT-Cre × Ai32 mice (n=4) (**Fig. 3A-B**). Following 1 week for retrograde transport, we identified EYFP+ cholinergic neurons throughout the rostral-caudal extent of the basal forebrain (**Fig. 3C, top**). Bead+ neurons clustered in two locations: a rostral-ventral-medial region that overlaps with the Horizontal Limb of the Diagonal Band and a second caudal-dorsal-lateral zone that falls within Nucleus Basalis (**Fig. 3C, middle**), where the highest density of ChACx cells were found (**Fig. 3C, bottom**) (Kim et al., 2016). Even in Basalis, the majority of ChAT+ neurons (~73%) did not contain retrobeads, underscoring the need for an approach that could functionally isolate if Basalis → ACx ChACx neurons from other neighboring ChAT+ neurons that do not project to ACx. This was accomplished by flashing blue light onto the surface of the ACx to trigger action potentials in cholinergic ACx axons and then recording the antidromic spike near the Basalis cell body (**Fig. 3D**).

**Fig. 3.**
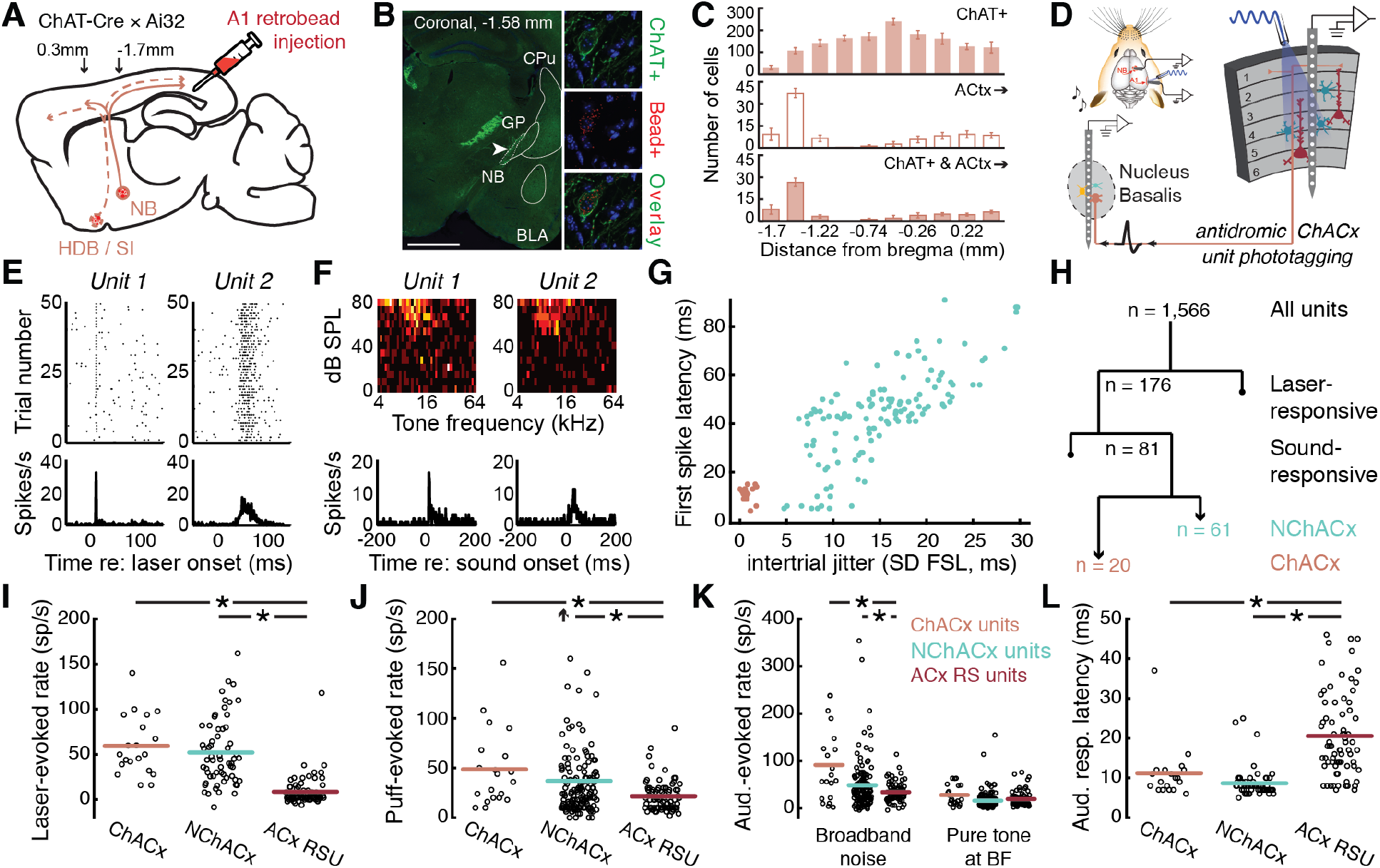
Multiplexed encoding of sensory and reinforcement signals in Nucleus Basalis. (***A***) Mid-sagittal mouse brain cartoon showing strategy for retrograde transport of fluorescent microbeads from ACx to cholinergic nuclei in the basal forebrain. NB = Nucleus Basalis. HDB / SI = Horizontal Limb of the Diagonal Band / Substantia Innominata. Arrows depict zone of anatomical reconstructions expressed relative to Bregma. (***B***) Coronal photomicrographs show EYFP fluorescence in cholinergic (ChAT+) neurons, microbeads in ACx-projection neurons and the overlay. CPu = caudate putamen; GP = globus pallidus; BLA = basolateral amygdala. Scale bar = 1mm. (***C***) Histograms of ChAT+, Bead+ or both ChAT+ and bead+ cells across the caudal to rostral extent of the basal forebrain. (***D***) Strategy for antidromic phototagging of Basalis ChAT+ neurons that project to the ACx (ChACx). (***E***) Rastergram (top) and PSTH (bottom) of laser-evoked spiking in two Basalis units. (***F***) Frequency response areas and tone-evoked PSTHs from the same units. (***G-H***) First spike latency (FSL) and FSL jitter evoked by the laser (*G*) in 81 tone- and laser-responsive Basalis units. Brown coloring = ChACx (n=20); Aqua coloring = indirectly activated neurons operationally defined as not ChACx (NChACx, n = 61). (***I-K***) Firing rates evoked by the ACx laser pulse (*I*), air puff (*J*) and either broadband noise bursts or BF tones (*K, left and right*) in ChACx, NChACx and ACx RSUs (n = 82). Upward arrow indicates one outlier. (***L***) First spike latency in ChACx, NChACx and ACx RS units evoked by broadband noise bursts. Horizontal lines represent the sample mean, each single unit is represented by a single point. Asterisks indicate significant (p < 0.05) post-hoc pairwise differences with the Kruskal-Wallis statistic. Statistical tests are adjusted for inflated false discovery using the Holm–Bonferroni correction for multiple comparisons.

In rare instances where brief (50 ms) photoactivation of ACx increased the firing rate of Basalis single units (n = 176 of 1,566 recorded units), we noted that responses fell into two categories: short-latency spikes with low jitter in spike timing (**Fig. 3E, unit 1**) or a longer latency burst of irregularly times spikes (**Fig. 3E, unit 2**). Either type of unit could have native auditory responses featuring well-defined tonal receptive fields with short-latency auditory responses (**Fig. 3F**). Among Basalis units driven both by laser and pure tones (n = 81), plotting first spike latency against the first spike latency jitter revealed a sparse set of Basalis units (1.2%, n =20 of 1,566 units) that stood apart from the remaining sample of Basalis recordings (**Fig. 3G**). We operationally defined the short-latency, low-jitter population as directly activated ChACx units, whereas the longer-latency irregular spikes presumably arose through synaptic transmission via reafferent or local circuits within Basalis and were therefore classified as Not ChACx units (NChACx) (**Fig. 3H**).

Optogenetic activation of cholinergic axon terminals increased the firing rates in Basalis units as well as ACx units, although cortical RSU activation (n = 82) was significantly weaker and presumably arose through intracortical disinhibition (Letzkus et al., 2011; Pi et al., 2013; Takesian et al., 2018) (Kruskal-Wallis, post-hoc pairwise comparisons p < 1 × 10^−6^, **Fig. 3I**). We confirmed an earlier report that basal forebrain units were driven by aversive reinforcement cues, as air puff-evoked spiking was significantly higher in either cell type than in ACx RSUs (Hangya et al., 2015; Harrison et al., 2016; Letzkus et al., 2011; Lin and Nicolelis, 2008) (Kruskal-Wallis, both pairwise comparisons p < 1 × 10^−6^, **Fig. 3J**). Importantly, both ChACx and NChACx units exhibited vigorous responses to noise bursts and pure tones before any conditioning had taken place (**Fig. 3K**). Unlike rostral basal forebrain units or classic descriptions of midbrain dopamine neurons, robust sound-evoked responses were observed in Basalis prior to conditioning (Lin and Nicolelis, 2008; Schultz et al., 1997). Auditory response latencies in Basalis units were even shorter than ACx RSUs, presumably reflecting monosynaptic inputs from medial, short-latency regions of the auditory thalamus (Chavez and Zaborszky, 2017; Hackett et al., 2011) (11.2 ±1.51 and 8.7 ±0.47 ms, respectively, vs. 20.5 ±1.16 ms; Kruskal-Wallis, both comparisons p < 0.02, **Fig. 3L**).

These experiments demonstrate that Basalis unit spike trains multiplex sensory and reinforcement cues. The short response latencies in ChACx and NChACx Basalis units would provide sufficient time for cholinergic inputs to engage the disinhibitory microcircuit in ACx ahead of the arrival of the afferent sensory trace (Letzkus et al., 2011; Pi et al., 2013; Urban-Ciecko et al., 2018). These findings support our hypothesis that ACx circuits could integrate CS and US inputs across a 5s gap because, in effect, there would not be a 5s gap separating the arrival of the auditory sensory trace and the transient ACh surge. But if Basalis →ACx units respond to all sounds, one might expect ACx receptive fields to reorganize even in response to sounds presented outside of a behavioral context. This was clearly not the case, as ACx receptive field plasticity was only observed in conditions where sounds were paired with reinforcement (i.e., CS+ conditioning, but not CS- alone) (Fig. 2I).

We reasoned that the influence of Basalis → ACx spikes must somehow be amplified during CS-US conditioning in a manner not observed when sounds are presented without behaviorally relevant consequences. To this point, we observed that ChACx units rapidly modify their tonal receptive fields to enhance frequencies near the CS+ frequency, much like ACx RSUs (**Fig. 4A**). The plasticity index for ChACx was significantly enhanced Early and Late in conditioning (one-sample t-test, p < 0.005) and was significantly greater for CS+ conditioning than CS- conditioning (n = 10/10 for CS+/CS- conditioning, p < 0.05). Unlike ACx receptive field plasticity, ChACx tuning reverted to baseline shortly after conditioning was terminated (Post-conditioning, one-sample t-test, p = 0.16), suggesting that plasticity within the cholinergic projection neurons might be involved in the induction, but not the maintenance of ACx plasticity (Chubykin et al., 2013). Importantly, enhanced responses to the CS+ frequency were significantly greater in ChACx units than NChACx units (n = 10/24, respectively, unpaired t-test, p < 0.001, **Fig. 4B**), suggesting that frequency-specific changes within the neuromodulatory input from Basalis might be limited to cholinergic cell types.

**Fig. 4.**
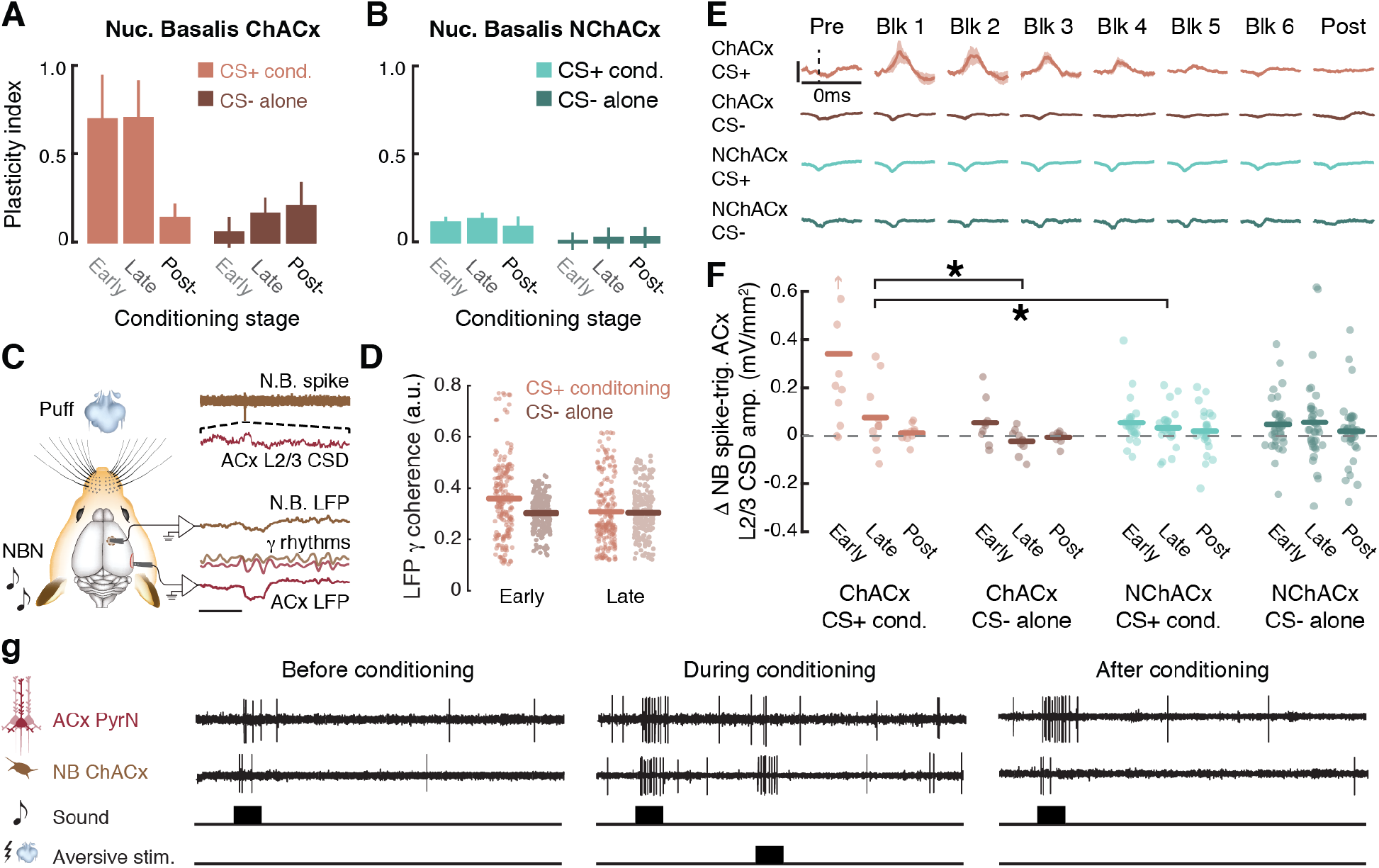
Local plasticity and enhanced feedforward cholinergic transmission from Basalis to ACx during auditory fear conditioning. (***A-B***) Plasticity index across conditioning stages for ChACx (*A*) and NChACx (*B*) units in CS+ conditioning (lighter hue, n =10/24 for ChACx/NChACx) and CS- alone conditioning (darker hue, n = 10/37 for ChACx/NChACx) measured during test blocks. (***C***) Simultaneous recordings of gamma-filtered local field potential signals (LFP, 40-80 Hz) and Basalis spike-triggered ACx L2/3 current source density (CSD) amplitude during conditioning blocks 1-3 (early) or 4-6 (late). (***D***) LFP gamma band coherence between ACx and Basalis during the early and late stages of CS+ delayed conditioning and CS- alone conditioning (176 CS+ and 198 CS- recordings. (***E***) Mean ± SEM Basalis spike-triggered ACx L2/3 CSD amplitude measured before, during and after each of six conditioning blocks. (***F***) Change in the Basalis-spike triggered L2/3 ACx CSD amplitude expressed relative to the pre-conditioning value. Each Basalis unit is represented by an open circle (upward arrow indicates one outlier). Solid horizontal bars indicate mean value. Asterisks indicate significant overall differences between CS+ delayed conditioning and CS- alone for ChACx units and in CS+ delayed conditioning blocks between ChACx and NChACx units (Wilcoxon Rank Sum tests, p < 0.05). Statistical tests are adjusted for inflated false discovery using the Holm–Bonferroni correction for multiple comparisons. (***G***) Illustration of proposed model for ACtx plasticity when long temporal gaps separate the auditory CS and an aversive US.

In addition to scaling up spiking rates at the CS+ frequency, we also observed a transient increase in the functional connectivity between Basalis and ACx at the initiation of learning. For this analysis, we turned to simultaneous electrophysiological recordings from Basalis and ACx during the NBN-air puff Conditioning blocks, rather than the interleaved pure tone Test blocks (Fig. 1c). Gamma rhythms in the local field potential (LFP) are well-established signatures of cognitive processing and inter-regional communication (Fries, 2015). LFP recordings from Basalis and ACx were analyzed during a 30s inter-trial interval period to avoid any potential influence of altered sensory-evoked responses (**Fig. 4C**). We found that gamma band (40-80 Hz) coherence between Basalis and ACtx was significantly increased during the initial blocks of CS+ conditioning compared to CS- alone blocks (n = 176/198 samples CS+/CS-, unpaired t-test, t=4.82, p < 5 × 10^−6^), but not during the second half of conditioning blocks (t = 0.46, p = 0.65, **Fig. 4D**). To directly estimate the “forward influence” of Basalis unit spikes on ACx, we measured changes in the amplitude of the ACx layer 2/3 current source density (CSD) signal that were temporally locked to spikes from individual Basalis units. Unsurprisingly, individual Basalis unit spikes were not associated with strong time-locked network activity from the upper layers of ACx during pre-conditioning baseline recordings. This weak overall influence was maintained throughout CS+ conditioning blocks in NChACx units and during CS- alone blocks for both Basalis cell types (**Fig. 4E**). Among ChACx units, however, we noted a striking increase in the strength of the feedforward spike-triggered cortical CSD in CS+ conditioning blocks that was significantly greater than what was observed in CS- alone blocks (Wilcoxon Rank Sum, p < 0.01) or in CS+ blocks from NChACx units (Wilcoxon Rank Sum, p = < 0.05; **Fig. 4F**).

## Discussion

Taken together, our findings demonstrate that electrophysiological signatures of brief sounds disappear within ~ 500 ms of stimulus offset in ACx, yet distributed forebrain circuits can bridge a CS-US gap ten times this long to support associative plasticity and learning (Fig. 2). This was likely not accomplished by maintaining an overt (Fig. S1) or covert persistent sensory trace that bridged the gap between the CS and US, because even an intracellular or spatially distributed sensory afferent trace would have been reinforced by the delayed ACh pairing experiment (Fig. 1F-G), in agreement with earlier findings (Metherate and Ashe, 1991). Rather, our experiments suggest that the 5s CS-US gap is effectively bypassed by scaling up the sensory-evoked firing rates and overall functional connectivity of cholinergic Basalis → ACx spikes in the initial stages of associative learning (**Fig. 4G**). These findings support the general model for a cortical plasticity arising from the convergent arrival of phasic ACh inputs and afferent sensory responses (Hangya et al., 2015; Letzkus et al., 2015; Weinberger, 2004), while challenging the specific assumptions that phasic cholinergic inputs from the basal forebrain are elicited only by the US and are invariant during learning.

The credit assignment problem addresses how causes are assigned to delayed effects. Beginning with Pavlov, the assumption has been that credit assignment during learning is solved by having the brain maintain a reverberating trace of the sensory CS throughout the delay interval until the reinforcement signal arrives (Pavlov, 1932). Here we find that rather than stretch out temporal trace of the CS until the US arrives, neuromodulatory cells that encode the US reach backward in time to increase their response to the CS and increase their functional connectivity with sensory brain areas that support long-term memory and adaptive behavioral changes.

## Acknowledgments

We thank K. Hancock for developing data collection software. We thank I. Carcea and A. Takesian for critical feedback on the manuscript;

## Author contributions

WG and DBP designed the experiments. WG collected and analyzed the data with supervisory input from DBP. DBP prepared the manuscript;

## Data and materials availability

Data and analysis code can be made available upon request to any researcher for purposes of reproducing or extending the analysis.

## Experimental Procedures

### Subjects

All procedures were approved by the Massachusetts Eye and Ear Infirmary Animal Care and Use Committee and followed the guidelines established by the National Institutes of Health for the care and use of laboratory animals. For freely moving fear conditioning experiments, 8 adult CBA/CaJ mice of either sex were used. For head-fixed fear conditioning experiments, 30 adult ChAT-IRES-Cre × Ai32 transgenic mice of either sex were used (Jackson Labs stock numbers 006410 and 012569, respectively).

### Preparation for awake head-fixed awake recordings

Mice were brought to a surgical plane of anesthesia with ketamine/xylazine (induction with 100 mg/kg ketamine and 10 mg/kg xylazine, with 50–60 mg/kg ketamine supplements as necessary). Buprivicaine was injected subcutaneously to numb the surgical site. Body temperature was maintained at 36.5° with a homeothermic blanket system (Fine Science Tools). The periosteum overlying the dorsal surface of the skull was completely removed. The skull surface was prepared with 70% ethanol and etchant (C&B Metabond). A custom titanium head plate was then cemented to the skull, centered on Bregma. After recovery, animals were housed individually. Animals were given at least 48 hours to acclimate to the head plate before any further experiments.

On the day of the first recording session, animals were briefly anesthetized with isoflurane (1.5% in oxygen) while a small craniotomy (0.5 × 1.0 mm, medial-lateral × rostral-caudal) was made along the caudal end of the right temporal ridge, 1mm rostral to the lambdoid suture to expose A1. A second craniotomy (1.0 × 1.0 mm, medial-lateral × rostral-caudal) was made centered at 2.5 mm lateral and 1.5 mm caudal to bregma also in the right cortex to access Nucleus Basalis. Small chambers were built around the craniotomies with UV-cured cement and filled with lubricating ointment (Paralub Vet Ointment). At the end of each recording session, the chamber was flushed, filled with fresh ointment, and capped with UV-cured cement (Flow-It ALC).

### Head-fixed awake recordings

On the day of recording, the head was immobilized by attaching the head plate to a rigid clamp (Altechna). Mice could walk freely on a disk that was mounted atop a low-friction silent rotor and a high-sensitivity optical rotary encoder. Continuous monitoring of the eye and disk rotation confirmed that all recordings were made in the awake condition. Recordings were performed inside a dimly lighted single-wall sound attenuating chamber (Acoustic Systems). For A1 columnar recordings, a single-shank linear silicon probe (NeuroNexus A1×16-100-177-3mm) was inserted into the auditory cortex perpendicular to the brain surface using a 3-D micromanipulator (Narishige) and a hydraulic microdrive (FHC) with the tip of the probe positioned approximately 1.3 mm below the brain surface, such that the top 2 electrode contacts were outside the brain, the bottom 2 contacts were in the white matter or hippocampus, and the middle 11-12 contacts spanned all six layers of the auditory cortex. At the beginning of the first recording session, several penetrations were made along the caudal-rostral extent of the craniotomy to locate the high-frequency reversal of the tonotopic gradient that demarcates the rostral boundary of mouse A1 *(29)*. For Basalis recordings, a second silicon probe (NeuroNexus A1X16-50-177-5mm) was inserted into the other craniotomy with the depth of the probe tip around 3.8 - 4.3 mm below the pial surface.

### Optogenetic and acoustic stimulation for neurophysiology recordings

Digital waveforms for the laser command signal and acoustic stimuli were generated with a 24-bit digital-to-analog converter (PXI, National Instruments) using custom MATLAB (MathWorks) and LabVIEW (National Instruments) scripts. Acoustic stimuli were presented via a freefield electrostatic speaker positioned 10cm from the left ear canal (Tucker-Davis Technologies). Stimuli were calibrated before recording using a wide-band ultrasonic acoustic sensor (Knowles Acoustics, model SPM0204UD5). The optical signal was generated with a calibrated 488 nm laser diode (LuxX, Omicron) coupled to an optic fiber. The fiber tip was positioned approximately 1cm above the exposed surface of A1.

With the silicon probe was positioned in an A1 column, we derived the laminar position of each electrode from the current source density (CSD) evoked by broadband noise bursts (50 ms duration, 4 ms onset/offset cosine ramps, 1 s inter-stimulus interval, 70 dB SPL, 100 repetitions). Frequency-rate functions from all recording sites were delineated from pure tone pips (250 ms duration, 4 ms onset/offset cosine ramps, 1 s ISI, 4—45 kHz with 0.1 octave steps, 70 dB SPL, 15 repetitions of each stimulus, pseudo-randomized). Frequency response areas (FRAs) were also measured using pure tone stimuli (250 ms duration, 4ms onset/offset cosine ramps, 1 s, ISI, 4 – 64 kHz with 0.1 octave steps, 0 – 80 dB SPL with 5 dB steps, 2 repetitions of each stimulus, pseudo-randomized).

### Single unit isolation and analysis of spiking activity

Raw signals were digitized at 32-bit, 24.4 kHz (RZ5 BioAmp Processor; Tucker-Davis Technologies) and stored in binary format. To eliminate artifacts, the common mode signal (channel-averaged neural traces) was subtracted from all channels. In experiments where simultaneous recordings were made from probes in A1 and Nucleus Basalis, the common mode removal was performed independently for each probe. Electrical signals were notch filtered at 60Hz, then band-pass filtered (300-3000 Hz, second order Butterworth filters), from which the multiunit activity (MUA) was extracted as negative deflections in the electrical trace with an amplitude exceeding 4 s.d. of the baseline hash. Single units were separated from MUA using a wavelet-based spike sorting package (wave_clus). Single unit isolation was confirmed based on the inter-spike-interval histogram (less than 3% of the spikes in the 0-3 ms bins) and the consistency of the spike waveform (s.d of peak-to-trough delay of spikes within the cluster less than 200 μs). The average trough-to-peak delay from each single unit formed a bi-modal distribution, allowing us to separate regular-spiking putative pyramidal neuron waveforms (> 0.4 ms) from fast-spiking waveforms (< 0.4 ms) (Figure S1A).

Frequency response functions were measured at least twice before the first conditioning block, between each conditioning blocks, and at least 4 times after the last conditioning block (Fig. 1C). CS-specific changes in firing rate were quantified by first normalizing the response between the values 0 and 1, such that a value of 1.0 equaled the response at the best frequency. We then computed the average fold-change in normalized response at frequencies far away from the paired/conditioning frequency (> ± 0.25 octaves, blue shaded region in Fig. 1E) and subtracted this value from the average fold-change in spiking at or near the paired/conditioning frequency (≤ 0.25 octaves, red). The resultant Plasticity Index is a marker of associative plasticity, where values greater than 0 indicate disproportionately enhanced spiking at the CS frequency and/or reduced spiking away from the CS frequency.

During conditioning blocks, the center frequency of the narrowband noise (NBN) conditioning stimulus was manually set by evaluating frequency tuning functions on each responsive electrode and selecting a center frequency that was at least 0.5 octaves away from the BF of all recorded units. On CS+ trials, NBN (4th order Butterworth band-pass filter, 0.25 octave bandwidth, 2 s duration with 5 ms cosine ramp at stimulus onset and offset, 70 dB SPL) was followed by a 5s silent period and then an air puff. The block/session organization of CS- conditioning were the same as the CS+ conditioning, except that no air puff was delivered. For cholinergic pairing experiments, air puffs were replaced with photoactivation of ACh axon terminals in A1 by shining blue light to the brain surface (488 nm, 2 s duration, 50 mW). Two versions of ACh pairing protocols were used: a delayed ACh photoactivation, where laser onset occurred 5s after sound offset, and a simultaneous photoactivation experiment where the ACh photoactivation overlapped with the sound presentation. Each of six individual conditioning blocks contained 10 pairing trials separated by a 40s inter-trial interval.

### Antidromic identification of A1-projecting ChAT neurons

To identify ChAT+ neurons in Basalis that project to the auditory cortex (ChACx), brief laser pulses were delivered to the surface of the exposed auditory cortex (488 nm, 50 ms duration, 50 mW, 1 s inter-trial intervals, 50 trials). Unit responses from the NB recording sites were analyzed as peri-stimulus time histograms (PSTHs). According to the response profile, three types of units were identified: direct antidromically activated units which exhibited short latency (5 – 15 ms) and low inter-trial jitter (< 2 ms); indirect antidromically activated units which exhibited long latency (typically 30 – 60 ms) and high inter-trial jitter (> 2 ms); and non-antidromically activated units which did not show significant responses to laser. Laser responses were monitored periodically during the experiment and only units with consistent response profiles were analyzed.

### Analysis of local field potentials and current source density

To extract local field potentials (LFP), raw signals were notch filtered at 60 Hz and down-sampled to 1000 Hz. The current source density (CSD) was calculated as the second spatial derivative of the LFP signal. To eliminate potential artifacts introduced by impedance mismatching between channels, signals were spatially smoothed along the channels with a triangle filter (5-point Hanning window). Noise-evoked columnar CSD patterns were used to determine the location of the A1 recording channel. Two CSD signatures were used to identify L4: A brief current sink first occurs approximately 10 ms after the noise onset, which was used to determine the lower border of L4 (white arrow Fig. S1A). A tri-phasic CSD pattern (sink-source-sink from upper to lower channels) occurs between 20 ms and 50 ms, where the border between the upper sink and the source was used to define the upper boundary of L4.

Basalis spike-triggered auditory cortex (ACx) CSD was calculated from the inter-trial interval period of conditioning blocks. CSD amplitude from layers 2/3 was captured before and after each Basalis spike time to calculate an average waveform. The peak to trough difference of the averaged waveform is defined as its amplitude. Magnitude-squared coherence estimate (mscohere in Matlab) was used to analyze spectral coherence of the LFPs between A1 and Basalis. Because LFP waveform from the Basalis probe were similar across channels, these signals were averaged to serve as a single Basalis LFP trace. For each Basalis-ACx channel pair during a single conditioning trial, the LFP coherence was calculated using one 2-s long waveform prior to onset of the NBN CS stimulus. These single-trial coherence data were averaged for each conditioning block. The gamma coherence of any conditioning block is defined as the averaged coherence value in the gamma band (40 – 80 Hz).

### Head-fixed fear conditioning

Mice were acclimated to head-fixation for at least an hour before conditioning began. To quantify head-fixed freezing behavior, the rotary wheel encoder signal was downsampled to 1 kHz and smoothed with a 100-point Hanning window using a zero phase lag digital filter. Movement velocity was calculated as the first derivative of the wheel position signal. Freezing was operationally defined for individual time samples during conditioning blocks as a zero-velocity signal. We computed the percentage change in the fraction of time samples with freezing behavior between a 5s time epoch immediately prior to CS onset relative to a 1s time window occurring 3-4s after CS offset (i.e., during the trace period for CS+ conditioning blocks). The percentage change in freezing was averaged across all single trials within the session.

### Anesthetized extracellular cortical mapping

Mice were brought to a surgical plane of anesthesia with ketamine/xylazine (induction with 100 mg/kg ketamine and 10 mg/kg xylazine, with 50–60 mg/kg ketamine supplements as necessary). Buprivicaine was injected subcutaneously to numb the surgical site. During the course of recording, the core body temperature of the animal was maintained at 36.5°C with a homeothermic blanket system (Fine Science Tools). Using a scalpel, a 4 × 3mm (rostrocaudal × mediolateral) craniotomy was made in the right auditory cortex, approximately centered on a point 2.8 mm posterior and 4.4 mm lateral to bregma, and the dura mater was left intact. The brain surface was covered with high-viscosity silicon oil and photographed. Simultaneous recordings were made from the middle layers of the right auditory cortex (420–430 μm from pial surface) with 2–4 epoxylite-coated tungsten microelectrodes (FHC). The location of each recording site was manually marked on a high-resolution photograph of the brain surface.

Sound stimuli were generated with a 24-bit digital-to-analog converter (National Instruments model PXI-4461) delivered to the ear canal via acoustic assemblies consisting of two miniature dynamic earphones (CUI CDMG15008–03A) and an electret condenser microphone (Knowles FG-23339-PO7) coupled to a probe tube. Stimuli were calibrated at the tympanic membrane in each mouse before recording. Normal function of the auditory periphery and accurate placement of the probe tube were confirmed by monitoring the threshold and amplitude of cochlear distortion product otoacoustic emissions. Frequency response areas were measured with pseudorandomly presented tone pips (50 ms duration, 4 ms raised cosine onset/offset ramps, 0.5–1 s inter-trial interval) of variable frequency (4–64 kHz in 0.1 octave increments) and level (0–60 dB SPL in 5 dB increments). A total of 533 unique frequency-level combinations were presented once or twice for a given recording site. Recordings were performed inside a double-wall sound attenuating chamber (ETS-Lindgren).

Raw neural signals were digitized at 32-bit, 24.4 kHz (RZ5 BioAmp Processor; Tucker-Davis Technologies) and stored in binary format. Subsequent analyses were performed in MATLAB (MathWorks). The signals were notch filtered at 60 Hz and then band-pass filtered at 400–3000 Hz with a second-order Butterworth filter. MUA spiking was detected as negative threshold-crossing events (−4.5 SDs from the mean). For each recording site, an FRA was constructed based on the onset portion of the tone-evoked MUA responses (10 – 40 ms), and the best frequency (BF) assigned to the frequency with the highest firing rate. As described previously, the primary auditory cortex (A1) and the anterior auditory field (AAF) were identified based on the stereotypical low-to-high-to-low frequency tonotopic organization across the caudal-rostral axis *(29)*. Tonotopy was reconstructed from 40-60 approximately evenly spaced individual recording sites with a 0.5 × 1.5 mm (lateral-medial × rostral-caudal) strip spanning the center portion of the A1-AAF extent. The high-frequency mirror reversal boundary separating A1-AAF was used as a common anchor point to align maps from different mice.

### Fear conditioning behavior – freely moving mice

Fear conditioning was performed inside an acoustically transparent enclosure (20 × 15 × 30 cm, L × W × H) resting atop electrified flooring (8 pole scrambled shocker, Coulbourn Instruments). The acoustic enclosure was maintained inside a double-wall sound attenuating chamber (ETS-Lindgren). Acoustic stimuli and foot shock trigger signals were generated on a National Instruments PXI system using custom software programmed in LabVIEW. Auditory stimuli were delivered through a calibrated electrostatic free-field speaker positioned above the apparatus (Tucker-Davis Technologies). Mice were given at least 5 min to acclimate to the apparatus before each conditioning session. The CS+ stimulus was a 2s long NBN burst centered at 16 kHz. The CS- stimulus was an NBN with 8 kHz as its center frequency. Each session consists of 6 alternating conditioning blocks between CS+ and CS-, with 10 trials in each block. During every CS+ trial, a foot shock (0.1 mA, 1 s duration) was delivered 5 s after the offset of the sound. Foot shocks were not delivered on CS- trials. Inter-trial intervals were randomly chosen between 30 s to 40 s. A passive exposure control group was presented with the same NBN stimuli without any foot shocks.

Following five consecutive days of conditioning, mice were placed in a novel context on Day 6 and given at least 5 min to acclimate to the apparatus before the testing. On test sessions, 20 trials of randomized CS+ and CS- were delivered without foot shocks (inter-trials intervals randomized between 60-90 s. Mouse movement were captured with a commercial webcam at 20 frames/s. Significant motion pixels were identified for each frame as pixels where values (16-bit) changed by more than 50 grayscale units. For each individual video frame, freezing was operationally defined in those containing fewer than 20 significant motion pixels. Percent time spent frozen was calculated for a 5 s period beginning at stimulus onset.

